# Perceptual consistency in phoneme categorization is driven by neural consistency and predicts improved speech-in-noise performance

**DOI:** 10.64898/2026.07.02.736174

**Authors:** Rose Rizzi, Jack R. Stirn, Zara Eisenhut, Gavin M. Bidelman

## Abstract

Listeners discretize the speech signal by assigning sounds to phonetic categories, though there is variability in how individuals accomplish categorization. Having more consistent categorization of sounds may be advantageous for understanding speech-in-noise (SIN). Though, it is unclear how different levels of neural processing in the auditory system reflect these perceptual differences. We recorded brainstem frequency-following responses (FFRs) and cortical event-related potentials (ERPs) while listeners actively labeled vowels along an acoustic-phonetic continuum using a visual analog scale. We computed intertrial consistency of neural responses to index the stability of listeners’ neural speech representations across stimulus presentations. We also assessed how faithfully midbrain and cortical responses represented stimulus acoustics using representational dissimilarity matrices (RDMs) computed across all token pairs. Neural RDMs were then compared with acoustic and phonetic category RDMs to assess whether FFRs and ERPs carried gradient vs. categorical information of the speech signal. We found greater behavioral consistency during phoneme labeling was correlated with improved SIN scores. Neurally, we found greater cortical or subcortical consistency predicted greater behavioral consistency. RDMs revealed subcortical responses retained more acoustic details, while cortical responses more closely reflected abstract phoneme categories. Our findings reveal important benefits of perceptual consistency to other domains of speech perception. We find perceptual consistency is driven by more consistent encoding of speech at either a cortical or subcortical level. More consistent sensory processing could provide a more stable readout of the speech signal to higher cortical brain areas which could confer advantages to later perceptual processes downstream.

## 1. Introduction

Listeners differ in how they accomplish phonetic categorization. In particular, they can vary in the style of their perceptual responses spanning from gradient to categorical modes of listening. Listeners have simultaneous access to both within- and between-category (or phonemic and subphonemic) information (Clayards et al., 2008; Massaro & Cohen, 1983; McMurray et al., 2008; Pisoni & Tash, 1974), but how they use this information to carve out their perceptual space varies. More gradient listeners utilize detailed acoustic information to inform their perception while more discrete listeners rely more heavily on abstract, category representations (Kapnoula et al., 2017; Rizzi & Bidelman, 2024). Thus, more discrete listeners have perception that is more strongly warped onto category labels and more gradient listeners maintain a more linear mapping of acoustics to the perceptual space (Kapnoula et al., 2017; McMurray, 2022; Rizzi & Bidelman, 2024). A listener’s categorization listening strategy (i.e., degree of gradience) can be assessed by measuring the slope of their identification curve from labeling phonemes along an acoustic-phonetic continuum similar to classic categorical perception experiments (Kapnoula & McMurray, 2021; Kapnoula et al., 2017; Kutlu et al., 2022; Myers et al., 2024; Rizzi & Bidelman, 2024; Wong et al., 2024). A shallower identification curve is representative of a more gradient listener, while a more discrete listener has a sharper identification curve.

Beyond individual differences in perceptual gradience/categoricity, listeners can differ in how consistently they assign speech sounds to categories (Kim et al., 2025c; McMurray, 2022; Myers et al., 2024). Some listeners have more stable (consistent) phonetic categorization, labeling the same tokens from a continuum similarly each time they are heard. Others have noisier (inconsistent) categorization, labeling identical tokens with different phonetic labels across multiple presentations. Early literature on categorical perception conflates the two constructs of consistency and gradience by using a two-alternative forced choice (2AFC) task (Apfelbaum et al., 2022; Kapnoula et al., 2017; McMurray, 2022). Because this task forces a binary decision rather than allowing graded responses, it is impossible to know whether a steep slope arises from consistent or discrete categorization and whether a shallow slope arises from noisy or gradient categorization (Apfelbaum et al., 2022). Visual analog scaling (VAS) paradigms have been adopted as better measures of individual differences in categorization (Apfelbaum et al., 2022; Kapnoula et al., 2021; Kapnoula et al., 2017; Massaro & Cohen, 1983; Rizzi & Bidelman, 2024, 2025). A VAS task allows graded responses, increasing variance in phoneme labeling across stimulus presentations. This task promotes more gradience, allowing more subtlety in responses than a forced choice task (Apfelbaum et al., 2022). Traditional identification curves and their slopes can still be derived to index gradience, but more notably, a VAS task provides a straightforward measure of consistency (1 – standard deviation of VAS responses per token) (Kapnoula et al., 2017; Kim et al., 2025c; Rizzi & Bidelman, 2025). Though there are mixed reports regarding whether these two perceptual indices are entirely independent (Honda et al., 2024b; Kapnoula et al., 2017; Kim et al., 2025b; Rizzi & Bidelman, 2024, 2025), there is increasing evidence that perceptual consistency might be a more sensitive and stable measure of categorization than gradience (Kim et al., 2025c; Myers et al., 2024; Rizzi & Bidelman, 2025).

There are several perceptual advantages for perceiving speech in a gradient and/or consistent manner. Gradiency has been linked to improved recovery from ambiguity (Kapnoula et al., 2021; McMurray et al., 2009) and perceptual flexibility (Kapnoula et al., 2017). Likewise, less consistent listening is associated with several speech-language disorders including dyslexia and developmental language disorder (DLD) (Kim et al., 2025a; Kim et al., 2025b; Kim et al., 2024). Importantly, the perceptual advantages of gradience and consistency may extend beyond the processing of isolated phonemes and transfer to more complex auditory processing like speech-in-noise (SIN) perception (Kapnoula et al., 2017; Myers et al., 2024; Rizzi & Bidelman, 2024, 2025). In a recent study, we found that more gradient listening strategies were associated with improved SIN performance on the QuickSIN (Rizzi & Bidelman, 2025). Moreover, when controlling for cognitive working memory capacity, we found consistency in phonetic labeling was a more robust predictor of QuickSIN performance than perceptual gradience (Rizzi & Bidelman, 2025). Other independent work has similarly demonstrated that gradience and consistency independently predict SIN performance, but that consistency is a more robust predictor due to its higher stability across different phonetic contrasts (Myers et al., 2024). Relatedly, having more “distinct” phonological representations yields a percept that is more robust to interference by noise (Elbro, 1996). At the other extreme, people with dyslexia may have less distinct phonological representations (Elbro et al., 1994), which could underlie both their decreased categorization consistency (Kim et al., 2025b) and deficits in SIN perception (Calcus et al., 2016; Dole et al., 2012; Ziegler et al., 2009).

While individual differences in categorization have been reported behaviorally, less is known about how these behavioral differences are represented at the neural level and across different stages of the auditory system hierarchy. Auditory event-related potentials (ERPs) have provided a window into cortical speech coding. ERPs are composed of slow positive and negative waves representing the time-course of sound coding, from early waves reflecting exogenous stimulus properties (N1; Näätänen & Picton, 1987) to later waves reflecting endogenous processing, including speech discrimination (P2; Alain et al., 2010; Ben-David et al., 2011), attention (P3; Picton & Hillyard, 1974) and semantic context (N400; Kutas & Hillyard, 1984). ERPs reveal gradient information is maintained in the time course of cortical speech coding through 900 ms (Bidelman et al., 2013; Sarrett et al., 2020; Toscano et al., 2018; Toscano et al., 2010). Categories are fully formed in the cortical ERPs by ∼200 ms around the latency of the P2 wave, suggesting categories are represented pre-perceptually (Bidelman et al., 2013; Chang et al., 2010). Earlier waves of ERPs (N1 ∼100 ms) show categorical-like changes across acoustic-phonetic continua in more discrete listeners, but remain largely linear in gradient listeners (Kapnoula & McMurray, 2021). These findings imply that category representations of the speech signal (i.e., the acoustic-to-phonemic mapping) might emerge earlier in the auditory system, potentially even prior to cortex for more categorical listeners and later in time for more gradient listeners.

Whether even earlier speech coding at the level of the auditory *brainstem* reflects categorical information or strictly mirrors gradient stimulus properties is more equivocal. Frequency-following responses (FFRs) are speech-evoked potentials that reflect sustained phase-locked activity predominantly in auditory midbrain (Bidelman, 2015a; Bidelman & Momtaz, 2021; Chandrasekaran & Kraus, 2010; Gorina-Careta et al., 2021; Smith et al., 1975). FFRs mirror spectral and temporal properties of speech with such high fidelity that the eliciting stimulus can be identified from the sonified neural waveform (Bidelman, 2018a; Galbraith et al., 1995). The strength and consistency of the FFR reflect the robustness of early subcortical auditory encoding (Kraus et al., 2017; Krizman & Kraus, 2019). FFRs also scale with perceptual abilities including SIN (Anderson et al., 2013; Bidelman & Momtaz, 2021; Parbery-Clark et al., 2009; Song et al., 2011b; Yellamsetty & Bidelman, 2019) and pitch processing (Bidelman et al., 2011b; Krishnan et al., 2010; Krishnan et al., 2012a; Marmel et al., 2013; Reis et al., 2021). Experience-dependent enhancement of FFRs have been observed in highly skilled listeners including musicians (e.g., Bidelman et al., 2011b; Musacchia et al., 2007; Musacchia et al., 2008; Parbery-Clark et al., 2009; Parbery-Clark et al., 2011; cf. Whiteford et al., 2025) and tonal language speakers (Bidelman et al., 2011a; Krishnan et al., 2009; Krishnan et al., 2005).

Earlier work using FFRs demonstrated that brainstem responses did not reflect categorical-level information of speech sounds (Bidelman et al., 2013). However, these studies utilized strictly passive listening tasks. More recent work and sophisticated paradigms have shown that brainstem FFRs to acoustic-phonetic speech continua do in fact reflect listeners’ phonetic category labels when recorded under *active* phoneme labeling tasks (Carter & Bidelman, 2023; Rizzi & Bidelman, 2023). This finding suggests top-down processing—presumably via the corticofugal efferent system—might shape speech representations into a more abstract, perceptually relevant form even at early, subcortical levels of auditory processing. However, it is unclear how distinct levels of auditory processing in brainstem and cortex reflect categorical and gradient perceptual listening strategies and moreover, how such differential weighting might account for SIN abilities.

Representational similarity analysis (RSA) can be used to assess such differential weighting, i.e., how multivariate responses (whether acoustic, neural, or otherwise) map to relevant features, perceptual models, or abstract representations of interest (Kriegeskorte et al., 2008). RSA is conducted by computing the correspondence (e.g., correlation) between two matrices each reflecting the dissimilarity between all tokens in a set. As the input, such representational dissimilarity matrices (RDMs) can be constructed as a matrix of pairwise dissimilarity (1 – correlation) scores between neural responses to an acoustic-phonetic speech continuum (Beach et al., 2021; Bidelman et al., 2013; Chang et al., 2010; Ou & Yu, 2022). Using RSA, these matrices can then be compared to conceptual or measured acoustic or categorical RDMs to quantify how closely neural speech coding aligns with gradient and category representations (Beach et al., 2021; Ou & Yu, 2022). In lieu of comparing neural RDMs to phonetic category and acoustic RDMs directly, neural RDMs can also be compared to behavior directly using multidimensional scaling (MDS) (Bidelman et al., 2013; Chang et al., 2010; Shepard, 1980). Regardless of method variant, RDMs are a powerful way to directly quantify whether a listener’s EEG responses reflect acoustic vs. phonetic coding across a collective set of speech stimuli.

Using RDMs, Chang et al. (2010) constructed neural dissimilarity matrices from intracranial responses to a 13 token continuum of CVs. They then used MDS to demonstrate neural representations in auditory cortex (Heschl’s gyrus) closely mirrored behavioral discrimination and identification around 110-150 ms after stimulus onset. Using scalp-recorded EEG and MDS, Bidelman et al. (2013) found that passive FFRs and early cortical ERP deflections (Pa: 20-40 ms, P1: 40-60 ms, and N1: 90-110 ms) did not mirror categorization behavior. However, cortical activity at 165-185 ms (P2 wave) closely followed listeners’ pattern of perceptual categorization, as in Chang et al. (2010). To directly address whether cortical stimulus representations more closely represented phonemic or subphonemic information, Beach et al. (2021) compared MEG-derived cortical RDMs to “perceptual” and “ambiguity” RDMs. The perceptual RDM was based on listeners behavioral identification, reflecting stimulus category membership. The ambiguity RDM was based on how consistently people labeled tokens in a 2AFC task, presumably reflecting more subphonemic detail since less consistent labeling under a 2AFC task could imply greater use of within-category information. They examined correspondence of neural RDMs to perceptual and ambiguity RDMs in three time windows where support vector machine (SVM) classification decoded behavioral reports (165-354 ms, 429-529 ms, 569-680 ms). In time windows where SVM classification could reliably predict behavior from cortical responses, both subphonemic and phonemic information were represented with phonemic representations being more prevalent in the 165-354 ms window and subphonemic representations being more prevalent in the 569-680 ms window. These results suggest gradient representations are maintained longer in the time course of speech processing, while categorical representations emerge earlier, around the time-frame of the P2 wave, but diminish in further time periods.

To investigate how gradient representations in brainstem and cortex change with individual differences in perceptual gradience, Ou and Yu (2022) used RSA to relate acoustic RDMs, derived from the stimulus set, to cortical and subcortical RDMs, derived from the corresponding ERP and FFR responses, respectively. Listeners labeled phonemes along a continuum from /ba/-/pa/ followed by passive EEG recording. They found subcortical FFRs more closely mirrored stimulus acoustics in more gradient listeners (shallower identification slopes) whereas more discrete listeners (steeper identification slopes) had more distinct cortical ERP representations that diverged from stimulus acoustics between 50 to 250 ms. This suggests acoustic information is transformed into phonetic categories at the cortical level and moreover, that how listeners weight gradient vs. phonetic features of the speech signal might depend on different stages of auditory processing. However, there are several critical limitations of this study. First, they did not compare neural RDMs to categorical RDMs to assess the relative similarity to both discrete and gradient information. Second, they also assessed gradience using a 2AFC task, which is prone to conflating consistency and gradience (Apfelbaum et al., 2022; Kapnoula et al., 2017; McMurray, 2022). Third, EEG was recorded under a passive task, which may minimize subcortical categorical representations (Carter & Bidelman, 2023; Rizzi & Bidelman, 2023). Thus, it remains unclear whether (i) perceptual gradience and/or consistency in phonetic hearing independently relate to cortical and subcortical speech representations during active listening and (ii) whether speech-FFRs and ERPs themselves vary in neural gradience and/or consistency.

Theoretically, the construct of consistency in categorization could be related to how consistently the auditory nervous system encodes speech across repeated presentations of a given stimulus token. Measures of neural consistency (i.e., trial-to-trial repeatability) are thought to reflect the stability of evoked neural activity (Krizman & Kraus, 2019). Indeed, prior work links inconsistent brainstem speech encoding with poorer language and reading abilities (Hornickel & Kraus, 2013; Neef et al., 2017; White-Schwoch et al., 2015) which, in turn, has been linked to inconsistent categorization abilities (Kim et al., 2025b; Kim et al., 2024). A higher fidelity neural representation of speech throughout the auditory system could provide a clearer perceptual readout and a more distinct phonological representation (Elbro, 1996), manifesting as more consistent phonetic categorization.

Honda et al. (2024b) examined this premise at the *subcortical* level by assessing relations between FFR response consistency and categorization consistency. They did not find a significant correlation between neural and behavioral measures of consistency. However, consistency of *passive* FFRs recorded to the syllable /da/ were correlated with consistency of labeling native and non-native speech stimuli that were entirely unrelated to the FFR task. Thus, the task and neural measures were incongruent as the FFR stimulus was not used in the behavioral identification task and the FFRs were recorded passively. Since categorical information is only present in FFRs recorded during active speech perception tasks (Carter & Bidelman, 2023; Rizzi & Bidelman, 2023), novel FFR paradigms are needed to better assess the real-time correspondence between neural and behavioral consistency to identical speech stimuli. Honda et al. (2024b) also only examined neural consistency subcortically (i.e., FFRs). It is possible that consistency in physiological encoding varies across distinct levels of auditory processing (ERPs and/or FFRs) and that such stability in neural representation(s) influences how consistent listeners are in their perceptual acoustic-to-phonetic mapping.

The role of neural consistency at a *cortical* level has not been as heavily investigated as subcortical consistency. Increased cortical ERP consistency has been linked to bilingualism (Krizman et al., 2014), selective attention (Strait & Kraus, 2011; Strait et al., 2014; Strait et al., 2015), and improved rhythmic sequencing (Tierney et al., 2017). Because subcortical and cortical levels of the auditory system are bidirectionally linked via corticofugal efferent pathways and these feedback connections differ between listeners in an experience-dependent manner (Gao & Suga, 2000; Krishnan et al., 2012b), consistency in speech coding may scale with individual differences in perception across distinct levels of auditory processing (Krizman et al., 2014). It is possible that highly stable speech coding subcortically drives perceptual consistency, though it is also possible that higher-level cortical speech coding more closely mirrors perception. Given the documented relationships between neural consistency and improved perceptual performance across both levels of processing, we hypothesized that coordinated consistency in auditory cortex and subcortex would predict increased categorization consistency as measured behaviorally.

Here, we aimed to assess the weighting of categorical and gradient stimulus information and neural consistency across brainstem and cortical levels of auditory processing during active speech identification tasks. To that end, we recorded FFRs and ERPs to an acoustic-phonetic continuum of vowel tokens while listeners labeled phonemes. We investigated whether FFR and ERP response consistency predicted perceptual consistency to determine if stability in neural speech representations at brainstem vs. cortical levels accounts for more/less stability in the perceptual organization of speech. We hypothesized that brainstem speech coding would be more stable than cortical speech coding given that FFRs have higher test-retest reliability than ERPs (Bidelman et al., 2018; Hornickel et al., 2012; Song et al., 2011a) and neural phase-locking decreases at higher levels in the auditory system (Joris et al., 2004). We further hypothesized that enhanced neural consistency would drive enhanced perceptual consistency. To test how gradient vs. discrete speech representations are weighted along the auditory system, we compared neural RDMs, derived from FFRs/ERPs, to gradient and categorical representations modelled from acoustic and perceptual dissimilarity matrices. We hypothesized that FFRs would have more gradient representations due to their stimulus-mirroring properties, while ERPs would have more categorical representations, representing a more abstract phonetic code emerging further along the auditory pathway. Finally, we explored whether the similarity between neural and representational matrices correlated with individual differences in categorization and SIN perception to examine whether gradient information is weighted more heavily throughout the auditory system in more gradient listeners or listeners with better SIN abilities. We hypothesized that more gradient listeners would have more gradient neural representations at both levels, while more discrete listeners would have gradient neural representations at the subcortical level which transformed to more categorical representations at the cortical level, extending findings from Ou and Yu (2022).

## 2. Methods

There were *N* = 40 young adults (22.78 ± 4.6 years; 15 male, 25 female) with 16.33 ± 2.8 years of education and 8.38 ± 6.4 years of self-reported formal music training in our sample. All participants were monolingual English speakers, had normal hearing (≤25 dB HL; 0.25-8 kHz octave frequencies), and were largely right handed (74% ± 37% Edinburgh Handedness Inventory; Oldfield, 1971). Participants provided written informed consent in accordance with a protocol approved by the Institutional Review Board at Indiana University and were paid $15 an hour for their time.

### 2.1 Stimuli and tasks

#### QuickSIN

We administered the QuickSIN test prior to categorization experiments (Killion et al., 2004). Listeners heard lists of 6 sentences in 4-talker babble with a signal-to-noise ratio (SNR) decreasing in 5 dB steps from 25 dB to 0 dB with each sentence. They repeated back what they heard and the experimenter recorded correct keywords. The total number of correct keywords were subtracted from 25.5 to arrive at SNR-loss, the listener’s SNR threshold relative to clinical norms. Lower scores indicate better SIN performance. SNR-loss was averaged across lists to arrive at the final SIN score for each listener. We ran 2-4 lists as time permitted, resulting in test-retest reliability between 1 (4 lists) – 1.3 (2 lists) dB ("Quick-SIN - Instructions for Use," 2024). Notably, this test-retest is much smaller than the range of our listeners’ actual QuickSIN scores (range = 7 dB).

#### Phoneme categorization

For the phoneme categorization experiments, listeners labeled 5 synthetic vowels along an acoustic-phonetic continuum changing from /u/ to /a/ (Bidelman et al., 2020; Carter & Bidelman, 2023; Rizzi & Bidelman, 2024, 2025). Tokens were sampled equidistantly from the continuum changing first formant frequency (F1) from 430 Hz to 730 Hz and were otherwise identical with respect to F0 (150 Hz), F2 (1090 Hz), and F3 (2350 Hz). Tokens were 100 ms and were gated with 10 ms ramps.

Prior to the EEG experiment, listeners completed a “baseline” categorization task delivered in PsychoPy (v2023.1.1) (VAS_baseline_ task) as in Rizzi and Bidelman (2025). Listeners heard 30 presentations of each vowel (150 total trials) via Sennheiser HD 280 circumaural headphones at 75 dB SPL. There was a 500 ms interstimulus interval (ISI) between trials. On each trial, they reported what they heard using a visual analog scale (VAS) with endpoints labelled “oo” and “ah” (Apfelbaum et al., 2022; Kapnoula et al., 2017; Massaro & Cohen, 1983; Rizzi & Bidelman, 2024). Average VAS distance and reaction times were reported for each trial.

Listeners then completed phoneme categorization tasks during EEG recording. Listeners categorized phonemes under a VAS task (VAS_EEG_) and a 2AFC task (2AFC_EEG_) with the task varying trial-by-trial. For the VAS task, listeners clicked along a scale as in the baseline VAS labeling task. For the 2AFC task, listeners clicked a button on the screen to report the category. We were most interested in categorization under the VAS task, which promotes more gradient listening and provides better measures of individual differences (Apfelbaum et al., 2022; Kapnoula et al., 2017). However, we included the 2AFC task as a control to examine categorization and neural coding under a traditional task paradigm that promotes more discrete listening. To further promote either discrete or gradient listening, listeners were instructed to “listen for the details” of the speech sounds during the VAS task and to “listen for the category” of the speech sounds during the 2AFC task.

To achieve adequate trial counts needed for FFRs (i.e., several 1000), stimuli were presented in a clustered ISI paradigm (Bidelman, 2015b) (see **Fig. 1**). A single token was presented 15 times with a fast (10 ms) ISI to elicit the FFR followed by a slowed ISI (1100 ms) and a final presentation to elicit the ERP. A behavioral response was obtained following the final slowed stimulus presentation. A total of 140 ERP (and behavioral trials) and 2100 FFR trials were obtained per listener and token for each task condition.

**Figure 1.**
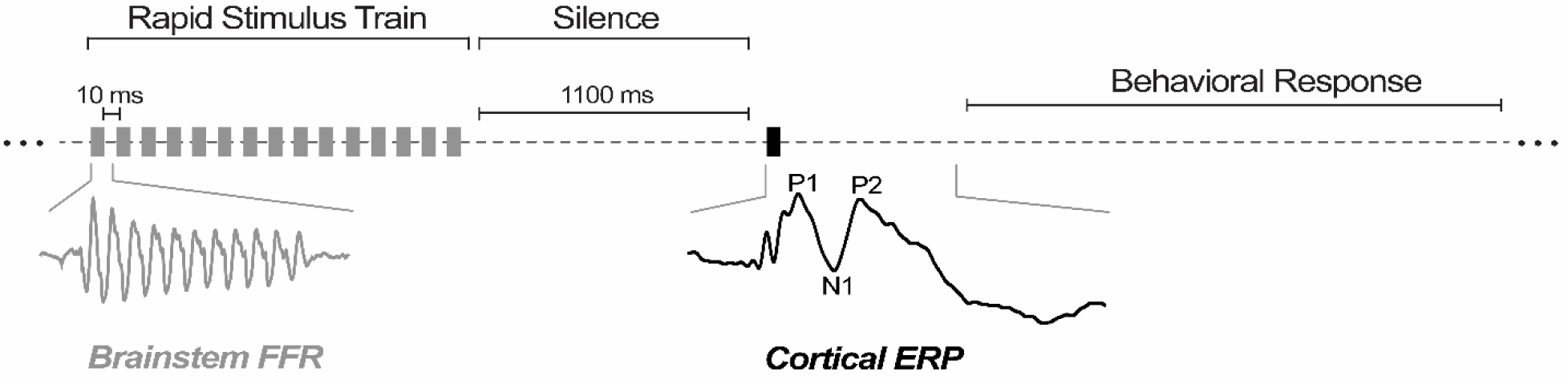
Stimulus presentation sequence recording FFRs/ERPs during active speech perception. Modified from Bidelman (2015b). Stimuli are presented in a train with a fast ISI (10 ms) to elicit the brainstem FFR, followed by 1100 ms of silence, then by a final presentation to elicit the cortical ERP. A behavioral response is made following the final slow presentation.

### 2.2 Behavioral data analysis

Individual subject identification curves were fit with a sigmoid P = 1/[1 + e^−β1(x−β0)^] and parameters were estimated using the *psignifit* function (Schütt et al., 2016) in MATLAB (v2024a). For each listener, the psychometric slope (β*1*) was estimated from VAS_baseline_, VAS_EEG_, and 2AFC_EEG_ responses to quantify perceptual gradience. Consistency (intertrial variance) was measured for each token and each listener in the VAS_baseline_ and VAS_EEG_ tasks as 1 – σ of VAS responses (distance clicked along the scale). Consistency cannot be reliably measured using a 2AFC task (Apfelbaum et al., 2022), so we did not measure consistency for 2AFC_EEG_. Reaction times (RTs) were calculated for each listener, token, and task type as the trimmed mean response latency across trials for a given condition. RTs <250 ms and >3000 ms were deemed fast guesses and lapses of attention and were removed from the analysis (Bidelman et al., 2013; Rizzi & Bidelman, 2025).

### 2.3 EEG recording and pre-processing

High density EEG was recorded using a 64-channel cap (QuikCap; Compumedics Neuroscan, Charlotte, NC) with Ag/AgCl electrodes located at 10-10 scalp positions (Oostenveld & Praamstra, 2001). EEGs were recorded using Neuroscan Curry 9 software and SynAmps RT Amplifiers (Compumedics Neuroscan, Charlotte, NC), were digitized at 5000 Hz, and were referenced to an electrode 1 cm posterior to Cz during recording. Recordings were later re-referenced to a common average reference offline. External electrodes on the outer canthi of the eyes and the superior and inferior orbit monitored eye movements. Principal component analysis was implemented in BESA Research 7.1 (BESA, GmbH) to spatially correct ocular artifacts (Picton et al., 2000; Wallstrom et al., 2004). Epochs >150 µV were rejected as artifacts and bad channels were interpolated based on visual inspection. EEGs were then separately bandpass filtered from 1-30 Hz or 100-1500 Hz (zero-phase filters, 48 dB/octave slope) to isolate cortical (ERP) vs. brainstem (FFR) activity, respectively. Filtered recordings were then epoched (-200-1000 ms for ERPs; -5-105 ms for FFRs), baseline corrected, and averaged across trials to generate ERPs and FFRs for each participant per token and task condition.

To resolve the intracranial generators underlying the scalp FFRs/ERPs, we transformed the multichannel sensory EEG data into source space using a virtual source montage with regional sources in brainstem and bilateral auditory cortex (Scherg et al., 2002). Source waveforms were obtained by matrix multiple of the sensor data by the inverse of the dipole leadfield matrix (forward volume conductor model) as described in Price and Bidelman (2021). We used the BESA default conductivities (S/m) of 0.33, 0.0042, 1.79, 0.33 for the scalp, skull, cerebrospinal fluid, and brain tissue compartments, respectively (Baumann et al., 1997). This digital re-montaging essentially applies a spatial filter to all electrodes (defined by the foci of our dipole configuration) to transform electrode recordings to a reduced set of source signals reflecting the neuronal current (in units nAm) as seen within each anatomical region of interest. This allowed us to calculate the weighted contribution of each source to the FFRs and ERPs thereby reducing each listener’s electrode recordings (64-channels) to 3 unmixed source waveforms reflecting activity in the midbrain for FFRs (Talairach coordinates: -0.064, -34.10, -12.15 mm) and bilateral auditory cortices for ERPs (Talairach coordinates: ±51.80, -22.53, +12.02 mm) (Scherg & Ebersole, 1994; Scherg et al., 2002). Activity from the y- and z-dipole orientations were averaged to describe the source FFRs, whereas the y-orientation (averaged across left and right hemispheres) was used to describe the source ERPs (Bidelman, 2018b; Price & Bidelman, 2021). These orientations optimally describe the direction of dipole current flow in the vertically oriented brainstem and tangential current flow in primary auditory cortex, respectively (Bidelman, 2015a, 2018b; Picton et al., 1999).

### 2.4 ERP and FFR neural consistency

We calculated cortical (ERP) and subcortical (FFR) neural consistency over the entire epoch window of each source waveform (FFRs: 0-105 ms; ERPs: 0-1000 ms) using a split-half, bootstrapped approach. For each participant, response trials to a given stimulus were randomly drawn, split in half (50/50%), and averaged. The two sub-averages were then Spearman correlated with one another, resulting in a measure of how consistently the FFR/ERP encodes the same stimulus across repeated trials. This procedure was then bootstrapped N=1000 times (with replacement) and repeated for each token (Fitzroy et al., 2015; Honda et al., 2024b; Krizman & Kraus, 2019; Otto-Meyer et al., 2018; Parbery-Clark et al., 2013; Tierney & Kraus, 2013; Tierney et al., 2017). To arrive at a measure of neural consistency comparable to the behavioral consistency measure, we subtracted the standard deviation of the bootstrapped correlations from 1 and z-scored the values to scale the measure across listeners.

Thus, a larger value represents a more consistent neural response.

### 2.5 Representational Dissimilarity Matrices (RDMs)

We constructed RDMs and used RSA to assess how similar neural responses were to (i) gradient features of the stimulus acoustics and (ii) phonetic category membership (Haxby et al., 2001; Kriegeskorte & Kievit, 2013; Kriegeskorte et al., 2008).

#### Neural RDMs

Only EEG data from the VAS task were used to ensure neural RDMs were from analogous conditions as the behavioral data. We computed dissimilarity between all pairwise EEG responses (separately for FFR and ERP response classes) to generate neural RDMs.

#### Stimulus RDMs

Similarly, we computed acoustic (i.e., gradient) RDMs via pairwise differences between the acoustic vowel waveforms. Dissimilarity was computed as 1-*r*_spearman_ for each token pair using the *pdist* function in MATLAB (Kriegeskorte & Kievit, 2013; Ou & Yu, 2022). The phonetic category membership RDM was constructed by setting matrix cells where tokens belong to the same category (i.e., tk1-2) to a dissimilarity of 0, and those belonging to opposite categories (i.e., tk1 and 5) to a dissimilarity of 1. The ambiguous midpoint of the continuum was set to a dissimilarity of 0.5 (see **Figure 9**). ^1^

We visualized the similarity between these stimulus model representational and EEG response matrices by computing pairwise Spearman correlations between cells (see **Fig. 9**). To provide a single measure to compare the stimulus and FFR/ERP RDMs, we computed the Euclidean distance between EEG and stimulus model RDMs with the diagonal removed to avoid artificially low distances (Bidelman et al., 2013; Chang et al., 2010; Guggenmos et al., 2018; Ritchie et al., 2017; Shepard, 1980). A larger distance between neural and acoustic RDMs indicates that the neural representation is more distant from (less similar to) the stimulus pattern in question (e.g., the acoustic or phonetic model comparison).

### 2.6 Statistical Analysis

Unless otherwise specified, we used mixed effects models (lme4 package version 1.1-32 in R version 4.2.1; Bates et al., 2015) to analyze differences in behavioral and neural dependent variables. Subjects served as a random effect where appropriate. Pairwise comparisons were Tukey-adjusted to account for multiple comparisons. Degrees of freedom for mixed models were estimated using Satterthwaite’s method. We also computed Spearman correlations between behavioral categorization measures and QuickSIN scores to assess relationships between SIN processing and gradience and categorization consistency.

## 3. Results

### 3.1 Behavioral data

To analyze whether gradience varied across our different task structures, we used a model predicting the slope of listeners’ phoneme identification functions from a fixed effect of task (3 levels: VAS_baseline_, 2AFC_EEG_, VAS_EEG_]) and random intercepts for subjects [i.e., slope ∼ task + (1|sub)]. We found a main effect of task [*F*(2, 76) = 87.23, *p* < 0.0001, 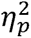 = 0.7]; slopes were expectedly steeper under 2AFC labeling than VAS labeling (2AFC vs. VAS tasks: *p* < 0.0001). There was not a difference between VAS_EEG_ and VAS_baseline_ labeling (*p* =0.98) (**Fig. 2A-C**). With few exceptions, slopes were correlated across tasks [2AFC vs. VAS_EEG_: *r*(37) = 0.42, *p* = 0.008; VAS_EEG_ vs. VAS_baseline_: *r*(37) = 0.40, *p* = 0.012; VAS_baseline_ vs. 2AFC: *r*(37) = 0.27, *p* = 0.098], suggesting perceptual gradience during phoneme labeling was relatively stable across different task structures.

**Figure 2.**
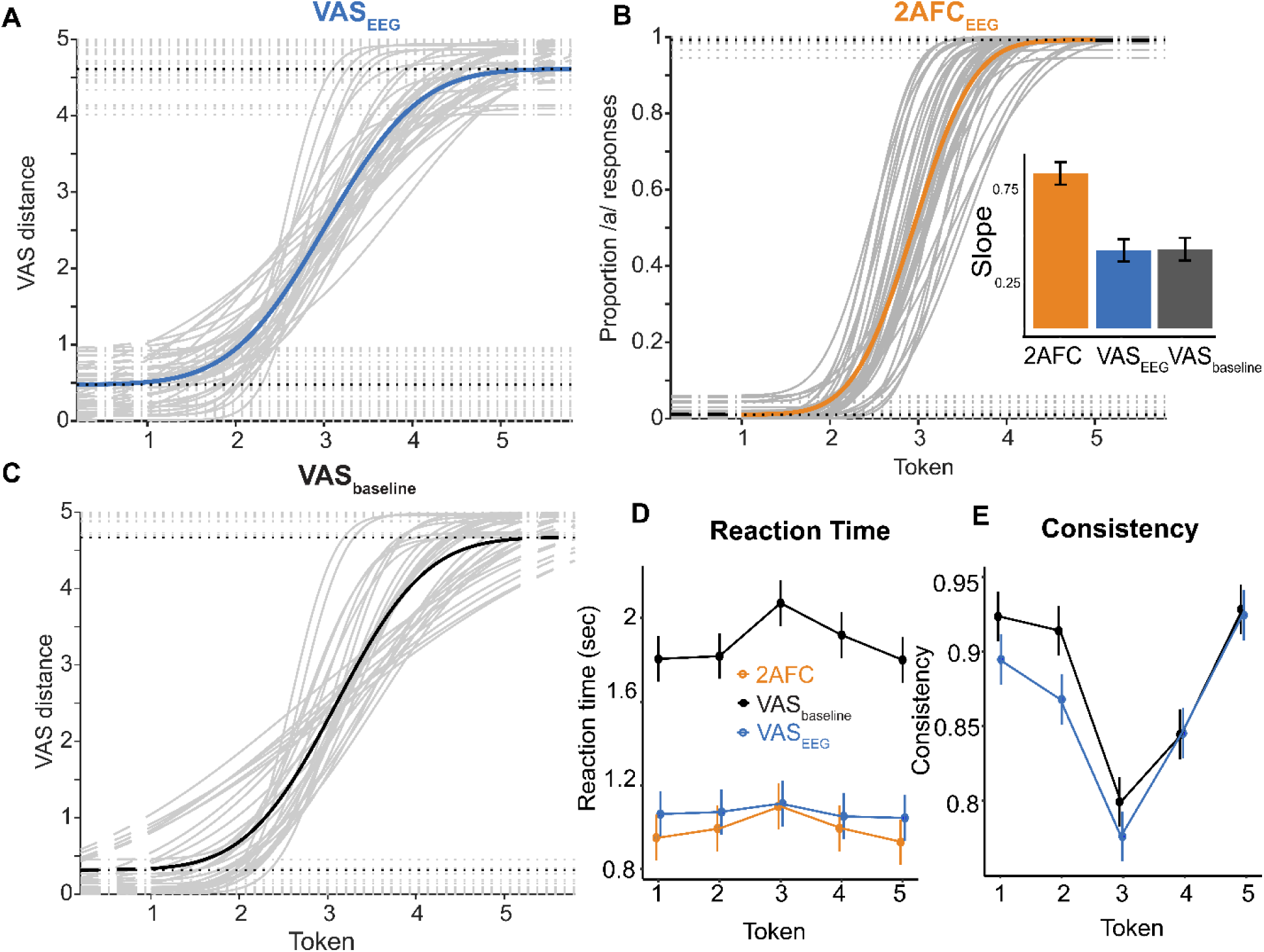
Behavioral phoneme labeling under the three task conditions. Identification curves (**A-C**) for different tasks show individual differences in gradience (dark grey traces = individual, colored traces = grand averages). Inset, slopes across tasks. Steepest slopes are observed in the 2AFC task; no differences in slope are observed in different iterations of the VAS task suggesting similar perception before and during EEG recording. (**D**) RTs across stimuli and tasks. RTs are slowest for the ambiguous token and in the VAS_baseline_ task. RTs are fastest in the 2AFC task. (**E**) Consistency across stimuli and tasks. Labeling is least consistent for the midpoint token and is less consistent in the VAS_EEG_ task. error bars = ± 1 s.e.m.

We next analyzed whether behavioral consistency and RTs varied across tokens [e.g., behavioral measure ∼ token * task + (1|sub)]. We found a main effect of token on labeling consistency [*F*(4, 351) = 96.95, *p* < 0.0001, 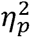 = 0.52] and RTs [*F*(4, 500) = 10.92, *p* < 0.0001, 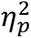 = 0.08] driven by less consistent and slower labeling for the ambiguous midpoint of the continuum relative to the endpoints [all Tukey-adjusted contrasts: (Tk1,2,4,5) vs. Tk3: *p* < 0.003] (**Fig. 2D**). There was also a main effect of task on labeling consistency [*F*(1, 351) = 15.87, *p* < 0.0001, 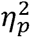 = 0.04] and RT speeds [*F*(2, 504) = 89647, *p* < 0.0001, 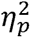 = 0.78]. RTs were slowest and labeling was most consistent in the VAS_baseline_ run (all contrasts *p* < 0.0001) (**Fig. 2E**). These results suggest the repeated stimulus train speeds RTs relative to the baseline task. There was no interaction between token and task on RTs (*p* > 0.06). In contrast, this interaction was significant for consistency [*F*(4, 351) = 2.81, *p* = 0.026, 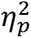 = 0.03], driven by less consistent token 2 labeling in the VAS_EEG_ task (*p* = 0.003). Listeners had least consistent labeling for the continuum midpoint across tasks, though listeners were less consistent overall in the prolonged VAS_EEG_ task (5x more trials).

We used the VAS_EEG_ behavioral data for subsequent comparisons with neural data since perceptual labeling occurred *during* EEG recording and thus provided a more veridical assessment of brain-behavior correlates.

We next calculated Spearman correlations between VAS_EEG_ categorization measures and QuickSIN scores (3 comparisons; Bonferroni-adjusted α = 0.0167). Importantly, we found QuickSIN scores were negatively correlated with categorization consistency (collapsed across tokens) [*r*(38) = -0.42, *p* = 0.007], indicating more consistent categorizers performed better on the SIN assessment (**Fig. 3**). We did not find a relationship between QuickSIN and phoneme gradience (slope) (*p* = 0.47) nor between QuickSIN and RTs (collapsed across tokens) (*p* = 0.07). These findings suggest the relationship between categorization skills and SIN processing are largely constrained to the construct of perceptual consistency (Rizzi & Bidelman, 2025) (cf. Myers et al., 2024; Rizzi & Bidelman, 2024).

**Figure 3.**
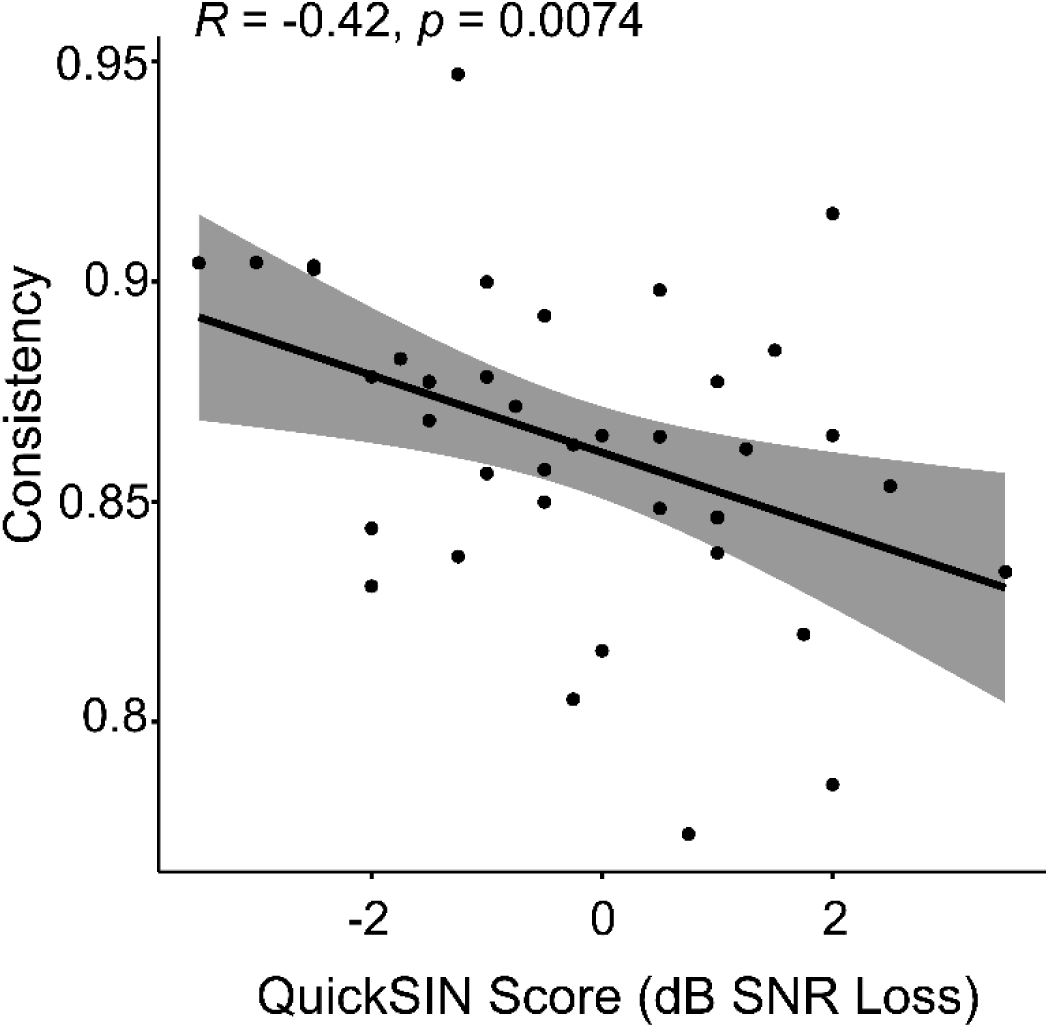
Categorization consistency is negatively correlated with QuickSIN scores. More consistent listeners had better (lower) SIN performance. Shading = 95% confidence interval.

### 3.2 Electrophysiological data

ERP (∼150 ms) and FFR scalp topographies for each token (collapsed across tasks) are shown in **Figure 4**. Both classes of response are maximal around fronto-central scalp locations (e.g., Cz/FCz), consistent with neural generators in the supratemporal plane (ERPs) and midbrain (FFR) (Bidelman, 2018b; Picton et al., 1999).

**Figure 4.**
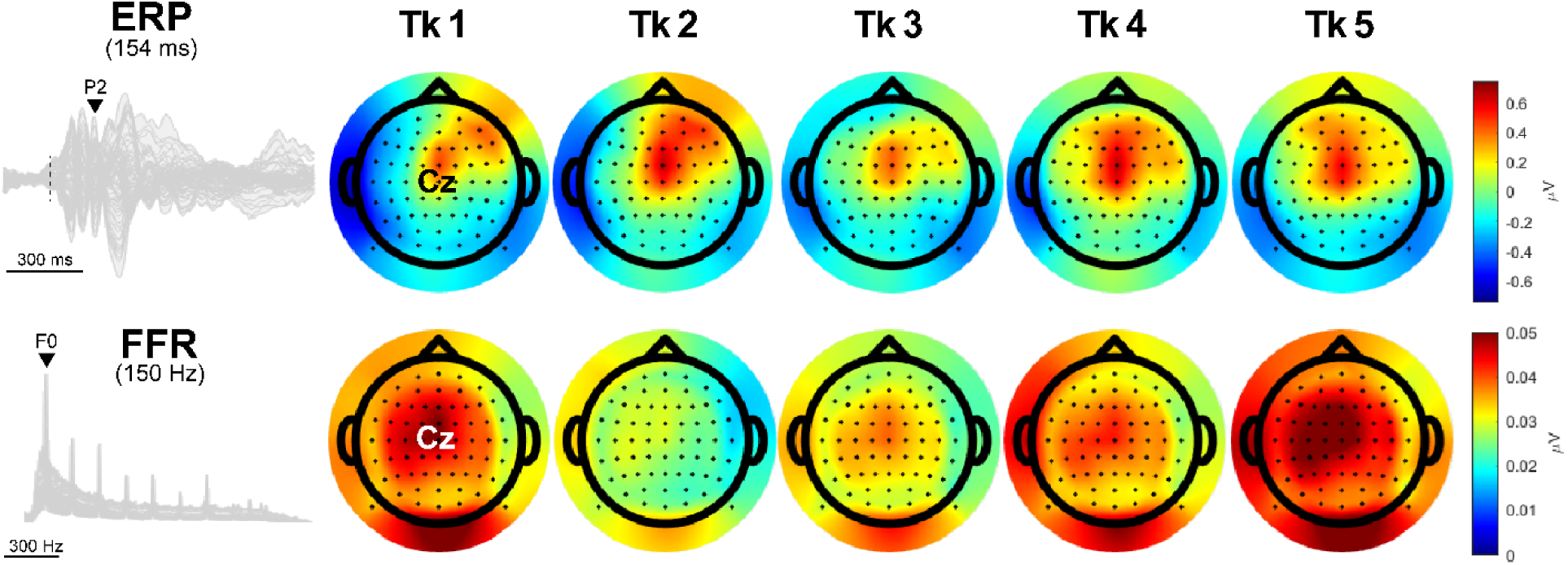
Topographies for ERPs and FFRs across tokens collapsed across tasks. ERP topographies are shown for the P2 (154 ms). Both FFRs and ERPs are maximal near the scalp vertex. FFR activity is stronger to endpoint tokens relative to others. ERP topographies similarly show weakest activity to the continuum midpoint relative to other tokens.

Grand average cortical ERP and FFR *sensor-level* waveforms and spectra are shown at FCz for all tokens and tasks in **Figure 5**. ERPs are shown common-average referenced and FFRs are mastoid referenced. ERPs for all tokens and tasks have similar morphology. FFRs mirror stimulus acoustics and track changes in F1 from 430 to 730 Hz. Amplitudes and latencies of individual ERP waves and FFR spectral amplitudes were not measured in favor of neural consistency measures and RSA.

**Figure 5.**
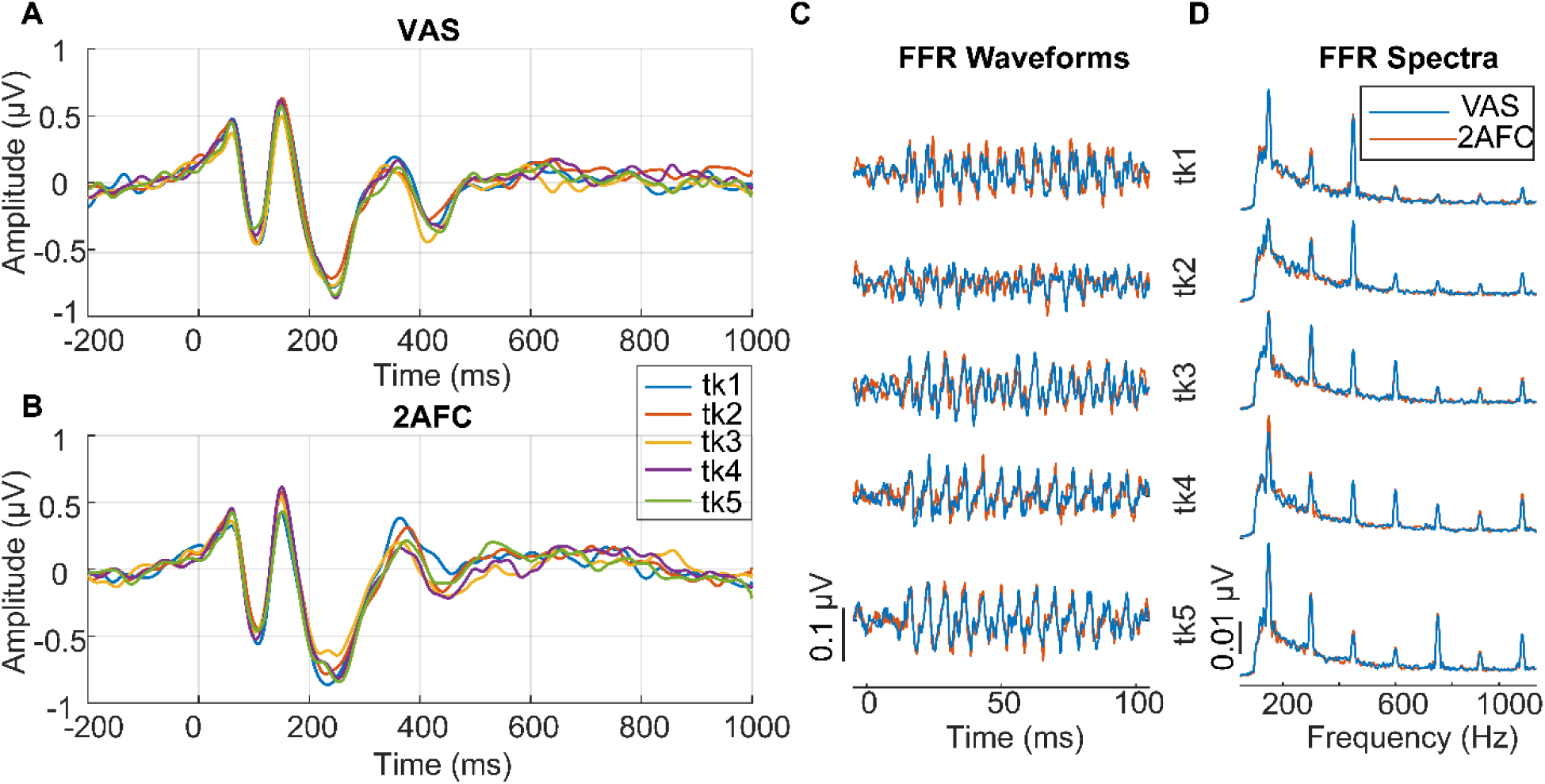
Cortical and brainstem speech-evoked potentials (FCz electrode) per speech token and task (VAS, 2AFC). **(A-B)** Grand average ERP traces. (**C-D**) Grand average FFR waveforms and their spectra.

Grand average *source-level* brainstem FFR and cortical ERP waveforms are shown in **Figure 6**. Source transformed FFRs (**Fig. 6A)** represent activity generated in the midbrain region. Likewise, canonical waves of the source ERPs (**Fig. 6B**) show activity estimated from left and right auditory cortex that closely resembles the scalp-recorded traces but parsed into contributions from each hemisphere (**Fig. 5A-B**).

**Figure 6.**
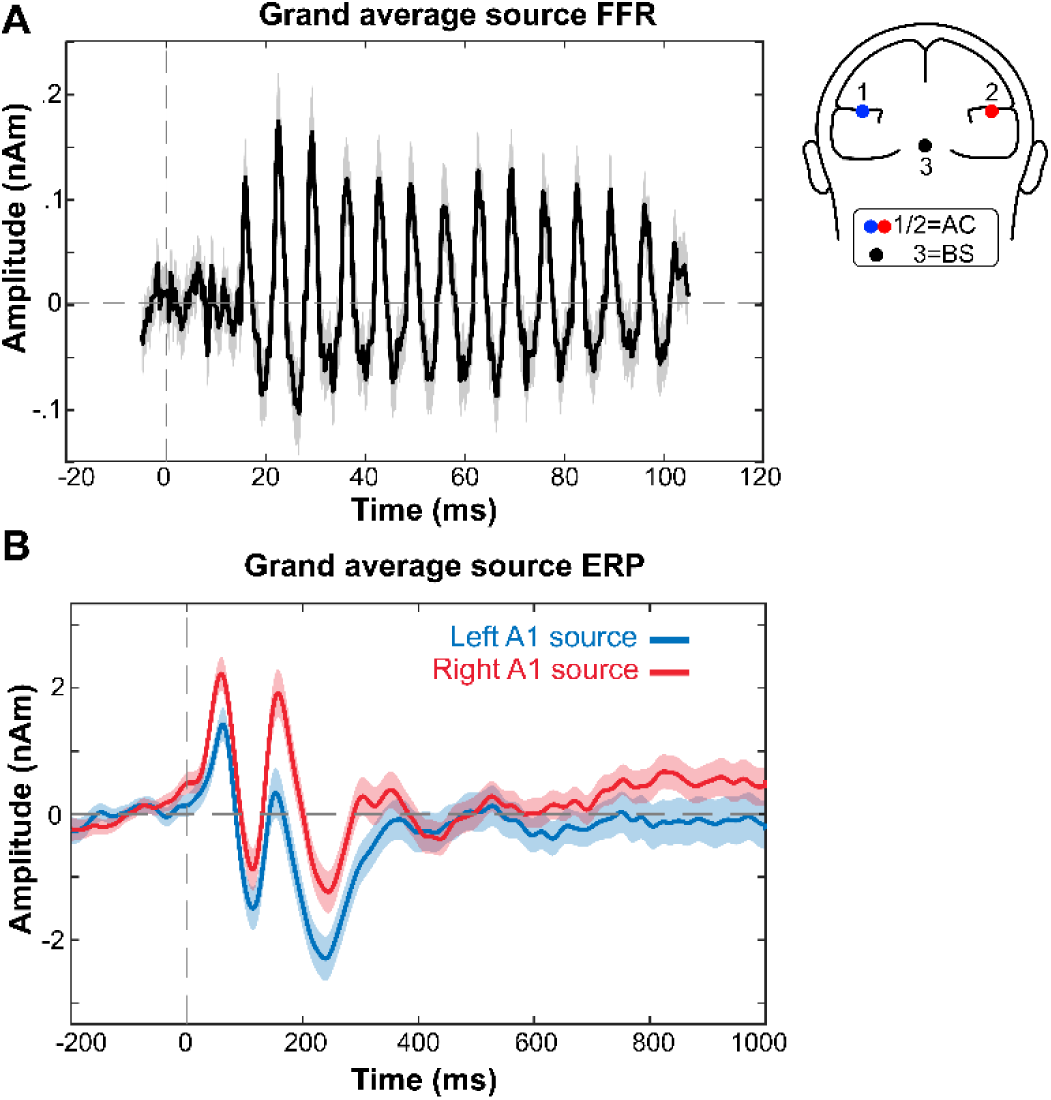
Brainstem FFR (**A**) and cortical ERP (**B**) grand average waveforms at source levels. Shading = 95% confidence interval. Waveforms are averaged across all listeners, task conditions, and speech tokens. Head inset, auditory cortical and brainstem dipole locations for source current waveforms (Talairach coord.: BS = -0.064, -34.10, -12.15 mm; AC = ±51.80, -22.53, +12.02 mm).

### 3.3 Neural consistency

To analyze differences in neural consistency, we used a mixed effects model with fixed effects of brain level (FFR vs. ERP), stimulus token, level*token interaction, and random intercept for subjects [i.e., neural consistency ∼ level*token + (1|sub)]. Neural consistency varied across levels of processing [*F*(1, 751) = 5.09, *p* = 0.024, 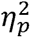 = 0.007] but with a level*token interaction [*F*(4, 751) = 8.22, *p* < 0.0001, 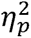 = 0.008]. The main effect of stimulus token was not significant by itself (*p* = 0.18). Tukey-adjusted multiple comparisons revealed that subcortical responses (FFRs) were more consistent than cortical responses (ERPs) overall. However, the level*token interaction was due to FFRs being notably more consistent than ERPs at tokens 1 and 3 (both pairwise *p* = 0.002) (**Fig. 7**).

**Figure 7.**
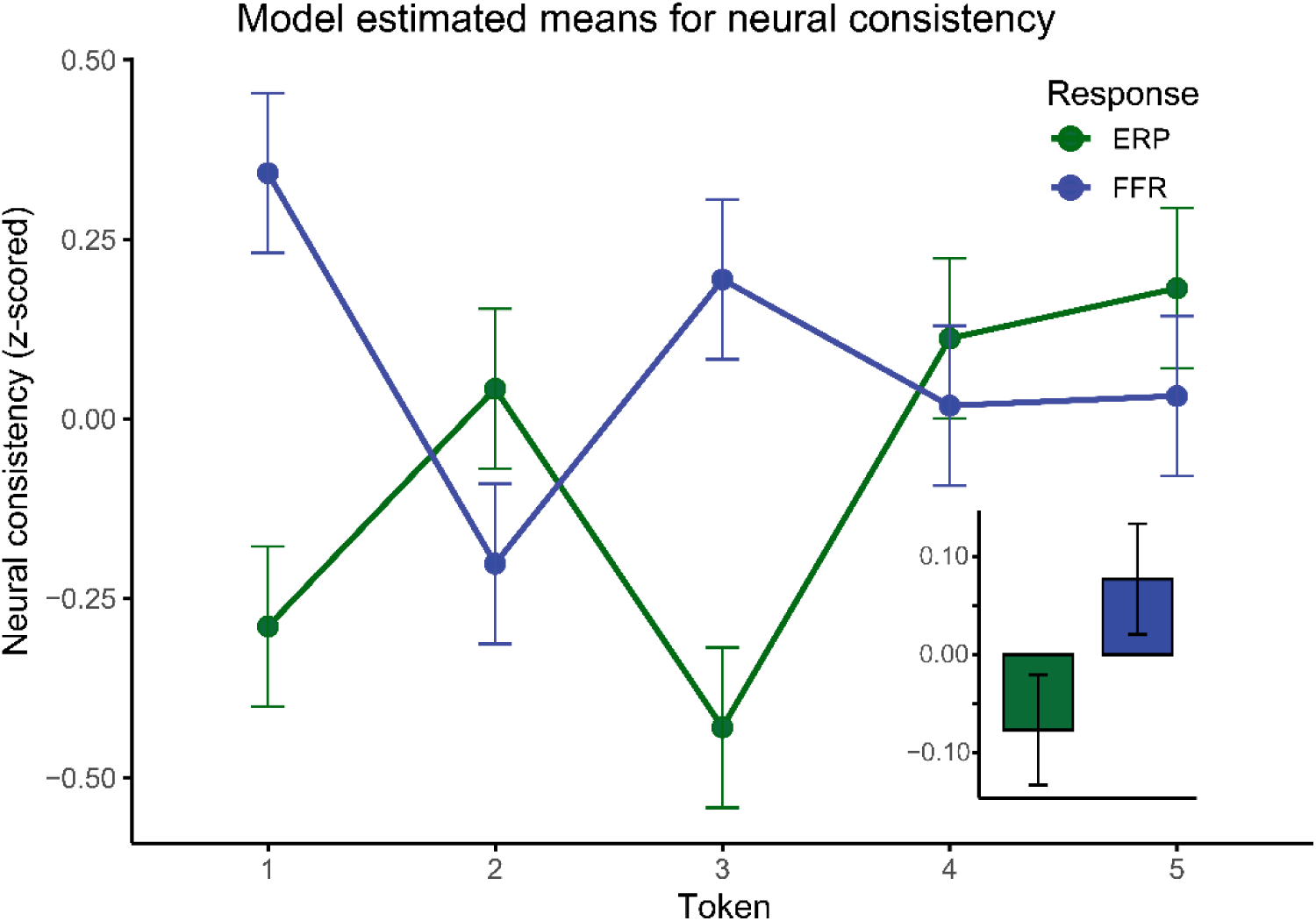
Neural consistency (z-scored) across stimuli and levels of processing. Data are collapsed across tasks. FFR and ERP consistency show nearly inverse patterns across the speech continuum. FFRs are more consistent overall than ERPs (inset) but the pattern of neural consistency across tokens interacts with anatomical level. error bars = ± 1 s.e.m.

To determine whether there was a correspondence between neural and behavioral responses, we modelled perceptual consistency (VAS_EEG_ task; Fig. 2E) with fixed effects of FFR and ERP consistency and their interaction [i.e., VAS_EEG_ consistency ∼ FFR consistency*ERP consistency + (1|sub)]. For this model, we only used EEG from the VAS task to assess brain-behavior relationships given those responses were collected under the same task and subject state. Both FFR consistency [*F*(1, 196) = 6.40, *p* = 0.012, 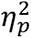 = 0.03] and ERP consistency [*F*(1, 165) = 7.64, *p* = 0.006, 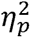 = 0.04] predicted behavioral consistency in phoneme labeling. However, and more critically, behavior was predicted by an FFR*ERP consistency interaction [*F*(1, 192) = 7.96, *p* = 0.005, 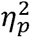 = 0.04]. Perceptual consistency was maximal when the neural consistency of ERPs and FFRs opposed one another. That is, trial-to-trial perceptual phoneme labeling reports were more stable when ERPs were consistent and FFRs inconsistent, or vice versa (**Fig. 8**).

**Figure 8.**
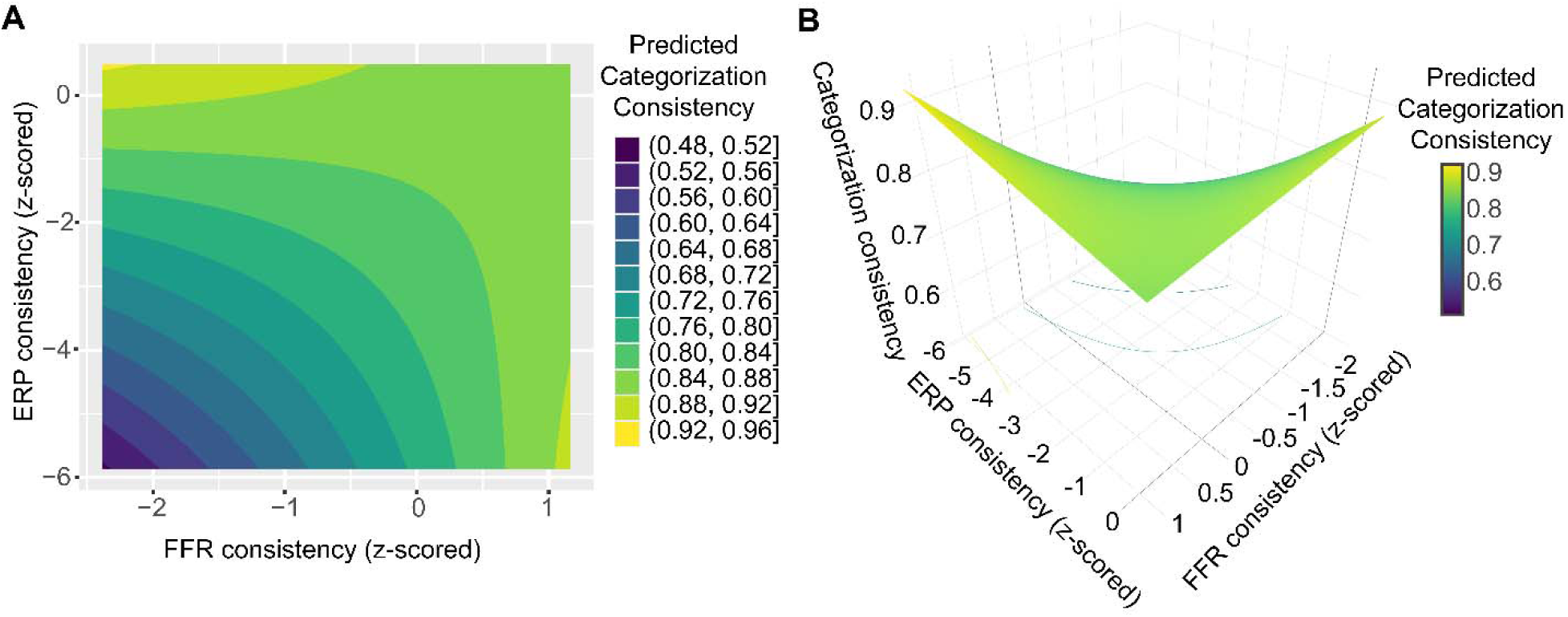
Neural ERP and FFR consistency interact to predict perceptual (categorization) consistency. (**A**) 2D grid illustrating the tertiary relation between ERPs, FFRs, and perception. Iso-contours (color coded bands) show model-predicted categorization consistency values. Yellow values = more consistent labeling behavior. (**B**) The same data projected onto a 3D surface highlighting the interaction between FFR and ERP consistency (saddle shape). Higher behavioral phoneme labeling consistency occurs when either ERP or FFR responses are highly consistent, and the other is not consistent. Poorest behavioral consistency occurs when both ERPs and FFRs are inconsistent.

### 3.4 RDMs

Finally, we assessed whether neural responses mapped to gradient vs. categorical representations of speech by comparing the distance between RDMs [i.e., neuro-acoustic distance ∼ level*representation type + (1|sub)]. Neuro-acoustic distance was strongly modulated by anatomical level [*F*(1, 105) = 15.67, *p* = 0.0001, 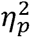 = 0.13], representation model type (gradient or categorical RDM) [*F*(1, 105) = 130.62, *p* < 0.0001, 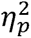 = 0.55], and, more critically, their interaction [*F*(1, 105) = 54.10, *p* < 0.0001, 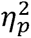 = 0.34]. The magnitude of distances revealed that FFRs more closely represented acoustic information than ERPs, while ERPs more closely represented categorical information than FFRs (**Fig. 9D**). This finding suggests that on the whole, brainstem FFRs encode more gradient features of speech than cortex, while cortical ERPs encode more phonetic-category information.

**Figure 9.**
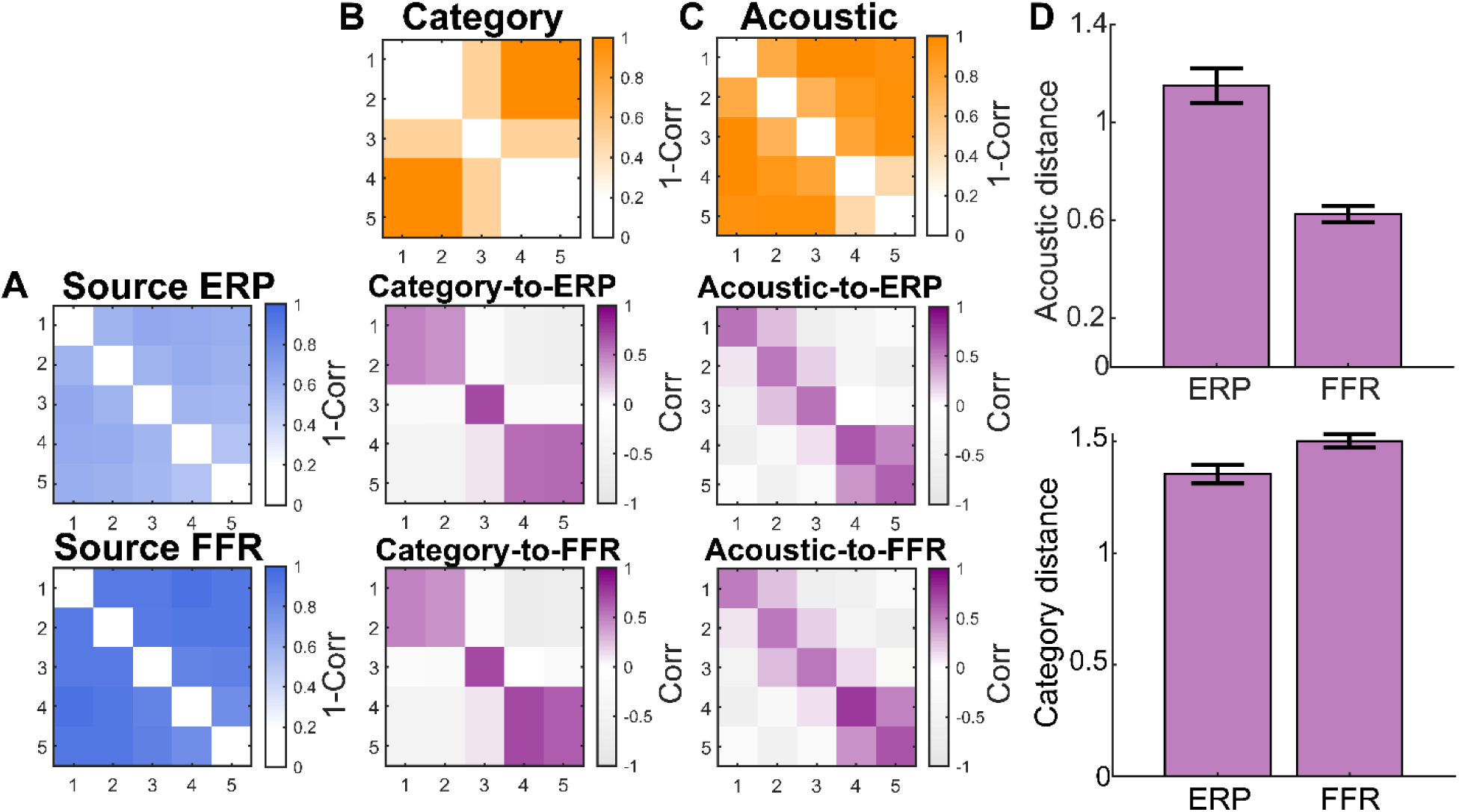
(**A**) Grand average neural RDM (pairwise 1 –r_Spearman_) for source-level ERPs (top) and FFRs (bottom). (**B**) Categorical RDM model representing phonetic categories (top); similarity (Spearman’s-r) between model and neural RDMs for FFRs and ERPs (middle, bottom). (**C**) Gradient RDM model representing acoustics. Otherwise as in B. (**D**) Average Euclidean distance between neural RDMs (panel A) and stimulus RDMs (panel B, C). FFRs have more gradient representations than ERPs, while ERPs have more categorical representations than FFRs. error bars = ± 1 s.e.m.

To assess whether these RDM effects mapped to behavior, we ran separate models with perceptual gradience (slope), consistency, and QuickSIN as the behavioral outcomes and FFR/ERP distance to the two respective RDM spaces as predictors [e.g., slope ∼ Dist_(FFR,acoustic)_ + Dist_(ERP,acoustic)_+ Dist_(FFR,category)_ + Dist_(ERP,category)_]. Since this model includes multiple distances derived from neural data which may not be independent, we assessed the variance inflation factor (VIF) of each predictor in the model to ensure results were not influenced by high multicollinearity. All VIFs remained under 1.2, indicating acceptable levels of multicollinearity. None of the RDM distance metrics predicted slope (all *p* > 0.209) nor consistency (all *p* > 0.213) in phoneme perception nor QuickSIN scores (all *p* > 0.126).

## 4. Discussion

We investigated how perceptual consistency and gradience in phoneme categorization are represented in speech-evoked potential across two primary stages of the auditory system. We simultaneously recorded source-resolved brainstem FFRs and cortical ERPs while listeners performed active speech labeling tasks. We further assessed the premise that consistency and/or gradiency in auditory perception might be related to more complex levels of linguistic analysis as indexed by speech-in-noise perception (Kapnoula et al., 2017; Myers et al., 2024; Rizzi & Bidelman, 2024, 2025). We found that listeners who labeled speech tokens more consistently performed better on the QuickSIN and had more consistent electrophysiological responses at brainstem (FFR) *or* cortical (ERP) levels (but not both stages). Using representational similarity analysis (RSA), we found that brainstem speech representations (indexed by FFRs) mapped more closely to stimulus acoustic features than phonetic categories. In contrast, cortical representations (indexed by ERPs) more closely resembled abstract phonetic category structure.

### 4.1 More consistent listeners have better SIN performance

At the behavioral level, categorization consistency (but notably not gradience) was related to SIN performance, supporting recent findings that more consistent listeners, who show highly repeatable labeling at the phoneme level, have superior SIN abilities (Myers et al., 2024; Rizzi & Bidelman, 2025). Though gradience has also been related to improved SIN comprehension (Bidelman et al., 2025; Myers et al., 2024; Rizzi & Bidelman, 2024), recent work suggests that perceptual consistency is more robust predictor of several perceptual processes [SIN: (Myers et al., 2024; Rizzi & Bidelman, 2025); reading and language skills: (Kim et al., 2025a; Kim et al., 2025b; Kim et al., 2024); second language learning (Fuhrmeister et al., 2023; Honda et al., 2024b)]. Consistency may be more trait-like than gradience as it is stable when assessed across a wide range of acoustic-phonetic continua (Kim et al., 2025c). While gradience in hearing can also persist across different stimulus materials (e.g., vowel vs. CVs) (Bidelman et al., 2025; Myers et al., 2024), it seems less stable, in a trait-like sense of perception (Kapnoula et al., 2021; Kim et al., 2025c; Myers et al., 2024). Increasingly, consistency has been posited as a more sensitive measure of categorization ability than gradience (slope) since it is assessed based on more subtle trial-to-trial variation in performance rather than the aggregate of many (Kim et al., 2025c).

Perceptual gradience, as measured in phoneme labeling tasks, might therefore be somewhat epiphenomenal, a byproduct of more variable responding that flattens the psychometric slope (Apfelbaum et al., 2022; Kapnoula et al., 2017; McMurray, 2022). Indeed, identification slopes measured in 2AFC tasks are related to consistency in VAS tasks, indicating that perceptual gradience is confounded by consistency when measured using conventional categorization methods (Honda et al., 2024a, 2024b; Kapnoula et al., 2017). Still, it remains equivocal in the literature whether gradience and consistency assessed via VAS tasks are truly independent. Some studies have found VAS slope and consistency are not correlated (Kapnoula et al., 2017; Rizzi & Bidelman, 2024), while others have found weak or inconsistent correlations across continua (Myers et al., 2024; Rizzi & Bidelman, 2025). While some work has investigated group differences among “canonically-categorical,” “categorical-but-noisy,” and “canonically-gradient” listeners (Kutlu et al., 2024). Here, we treated gradience and consistency continuously and in separate models to assess graded relationships with both measures across all listeners. This choice seems most pragmatic in light of the partial independence of perceptual gradience and consistency observed across studies.

Our finding that consistency but not gradience relates to SIN performance supports the notion that consistency is a more sensitive measure of individual differences and the linking factor relating categorization and SIN abilities. It is plausible that a relationship between consistency and SIN could have emerged in the opposite direction, implying decreased flexibility aids noise-degraded listening (Kapnoula et al., 2017). Instead, our hypothesis, bore out by data, suggests listeners with more consistent categorization perform better on the QuickSIN. This finding agrees with prior literature demonstrating listeners with poorer reading and language abilities have less consistent categorization (Kim et al., 2025a; Kim et al., 2025b), poorer SIN abilities (Chandrasekaran et al., 2009; Dole et al., 2012; Ziegler et al., 2009), and less robust neural speech coding (Chandrasekaran et al., 2009; Hornickel & Kraus, 2013; White-Schwoch et al., 2015). While several cognitive factors are also important predictors of SIN perception (e.g., Dryden et al., 2017; Hoover et al., 2017; Yoo & Bidelman, 2019), at the behavioral level, our findings underscore the importance of perceptual consistency as a key ingredient not only for phoneme-level categorization but also higher-levels of speech analysis (Myers et al., 2024; Rizzi & Bidelman, 2025).

### 4.2 Subcortical speech coding is more consistent than cortical speech coding

At the neural level, our hypothesis that brainstem responses would be more consistent overall than cortical responses was supported by our finding that FFR consistency was greater than ERP consistency. This finding mirrors Krizman et al. (2014) who found passively-evoked FFRs were more consistent than ERPs to a /da/ stimulus. FFRs are generally more stable and have higher test-retest reliability both between- and within-subjects than ERPs (Bidelman et al., 2018; Hornickel et al., 2012; Song et al., 2011a) which might account for such findings. The larger precision of phase-locked neural firing in auditory brainstem likely drives the enhanced repeatability and consistency of FFRs relative to ERPs (Krizman et al., 2014; Shadlen & Newsome, 1998). One could argue that the rapid stimulus sequencing of our clustered presentation might have biased FFRs to be more consistent. However, we find this account unlikely given that FFRs are still more repeatable than ERPs even under conventional stimulus paradigms using fixed rather the clustered ISI (Bidelman et al., 2018).

### 4.3 Neural consistency at either brainstem or cortical levels predicted perceptual consistency

Closely related to the present study, Honda et al. (2024b) assessed whether neural consistency of passively evoked FFRs (/da/ stimulus) was associated with categorization consistency of native and non-native phonetic speech contrasts but found no relationship between brainstem responses and behavior. However, a major limitation of that study was the use of strictly passive listening tasks which limits interpretation of the FFR data and understanding of how electrophysiological activity maps to perception. Importantly, categorical (i.e., phonetic) information is only present in the FFR under active listening conditions (Bidelman et al., 2013; Carter & Bidelman, 2023; Rizzi & Bidelman, 2023). Though it has been methodologically challenging in the FFR literature, our paradigm allowed us to successfully record brainstem responses during active speech categorization tasks in real time. This paradigm improves upon prior studies and allowed us to calculate neural consistency from FFRs while listeners simultaneously performed active phoneme labeling tasks. Thus, our neural and perceptual categorization consistency measures were derived under identical speech perception tasks, subject and attention states, and experimental demands. Further departing from Honda et al. (2024b), we also sought to examine the interplay of neural consistency in both brainstem and auditory cortex via concurrent recording of FFRs and ERPs and source analysis.

We originally hypothesized that coordinated consistency in brainstem and cortex would predict perceptual consistency. Instead, we found a critical interaction across levels in how these two classes of evoked responses modulated behavior. More consistent categorizers had either more consistent ERPs or FFRs, while less consistent listeners had inconsistent ERPs and FFRs. This interaction suggests that higher neural consistency in speech encoding is advantageous for perception only when it is present at *either* early or late stages of the auditory pathway, only partially supporting our hypothesis. Similarly, maturation of consistency in the auditory system differs across functional levels of processing with subcortical responses becoming less consistent (Skoe et al., 2015) and cortical responses becoming more consistent (i.e., less variable) with age (Fitzroy et al., 2015; Krizman et al., 2015). This inverse maturation pattern suggests that the *relative* stability of signal coding in different functional regions of the auditory brain hierarchy could be an important factor in auditory perception. Indeed, Centanni et al. (2022) found adults with dyslexia had decreased neural consistency during phoneme labeling in only one of six cortical regions associated with auditory perception, language, and reading (i.e., left supramarginal gyrus) rather than the entire auditory-linguistic circuit. This finding suggests increased neural consistency might not be homogenous across all auditory processing regions involved in decoding a task. Instead, specific brain areas might subserve perceptual stability rather than changes at the full brain level.

Top-down processing may influence the FFR (Galbraith et al., 2003; Price & Bidelman, 2021) but the effects of attention on brainstem potentials tend to be small and their existence is still debated (Galbraith & Kane, 1993; MacLean et al., 2024; Varghese et al., 2015). In contrast, cortical responses have much larger influences of attention (Alho, 1992; Galbraith & Kane, 1993; Näätänen & Teder, 1991), suggesting that ERPs are more susceptible to modulation by cognitive or attentional state on a trial-by-trial basis. Conversely, FFRs are a highly stable neuro-microphonic response with arguably less top-down influence than ERPs. Weaker top-down influences at brainstem vs. cortical stages of the auditory system might account for the higher consistency in neural representation we find in speech-FFRs vs. their ERP counterparts (Krizman & Kraus, 2019).

Our far-field recordings are also consistent with single-unit recordings showing a transformation in sound processing from brainstem to cortical levels. There is less redundancy in neuronal firing patterns in auditory cortex relative to subcortex, suggesting the neural code becomes more abstract higher along the central auditory pathway (Buck et al., 2025; Chechik et al., 2006; Tsunada et al., 2011). Neural representations from cochlear nucleus to auditory cortex become progressively decorrelated, corresponding to more distinct representations of sounds in the neural space along the ascending central auditory pathway (Gosselin et al., 2025). Similarly, we find subcortical FFRs are more consistent and robust in representing stimulus acoustic features of the speech signal. In contrast, less consistent responses in the ERPs could be in part due to the greater abstraction in the neural code observed at the cortical level.

Because neural representations are transformed along the auditory pathway (Buck et al., 2025; Chechik et al., 2006; Gosselin et al., 2025; Tsunada et al., 2011), it is likely that neural consistency differs across early and higher levels of processing. In addition to abstraction across the neural hierarchy, nonlinearities in auditory processing introduced along the afferent auditory system could magnify small variations in early responses (David & Shamma, 2013; Sharpee et al., 2011), potentially resulting in some listeners having consistent subcortical responses, but inconsistent cortical responses. Likewise, attention could increase stability of cortical representations despite noisy sensory inputs (Mesgarani & Chang, 2012; O’Sullivan et al., 2014), which could result in some listeners having consistent cortical responses but inconsistent subcortical responses. It is also possible that listeners weight subcortical and cortical speech coding differently, with some relying more on brainstem consistency and others relying more on cortical consistency. Thus, listeners who rely on consistency at a higher level of the auditory system (cortical consistency) may use more attention to drive their categorization behavior. In contrast, other listeners may rely on higher fidelity stimulus representations in early auditory processing stages (subcortical consistency) to make perception more automatic and drive consistent categorization.

More broadly, higher fidelity and more “distinct” neural representations of sound seem to facilitate downstream perceptual processing. For instance, more robust and distinct neural phonological representations may facilitate speech perception in noise (Bidelman et al., 2026; Elbro, 1996). Similarly, less robust subcortical encoding has been associated with poor speech processing in several neurodevelopmental disorders (e.g., dyslexia, autism) (Hornickel & Kraus, 2013; Neef et al., 2017; Otto-Meyer et al., 2018; White-Schwoch et al., 2015). Conversely, more consistent subcortical encoding has been reported in the FFRs of bilinguals (Krizman et al., 2014) and listeners with enhanced rhythmic ability (Tierney & Kraus, 2013; Tierney et al., 2017). Thus, it is plausible that consistent neural representations of speech provide a basis for perceptual consistency observed in our normal hearing listeners (Honda et al., 2024b) and, in turn, the impoverished or improved speech processing observed across different clinical and expert populations.

### 4.4 Gradient and categorical representations varied across processing levels

RDM analysis allowed us to assess the weighting of gradient and categorical information coded in EEG responses and thus whether brainstem and cortex code for lower vs. higher-level attributes of the speech signal (i.e., acoustic vs. phonetic representation). We found a double dissociation in how auditory brainstem and cortex coded continuous vs. discrete properties of speech sounds, supporting our hypothesis that gradience and categories would be more strongly represented subcortically and cortically, respectively. Euclidean distance revealed that FFRs were more similar to gradient acoustic representations than ERPs. In contrast, ERPs more closely resembled phonetic representations than FFRs. These results support Ou and Yu (2022) who similarly found subcortical RDMs more closely resembled stimulus acoustics than cortical RDMs. The small distance between FFRs and gradient acoustic RDMs in our data suggests that FFRs closely maintain within-category information. A more robust stimulus-mirroring property of the FFR naturally lends itself to encoding gradient acoustic features and may explain why category-relevant information is typically not observed at early, brainstem levels (Bidelman et al., 2013). Still, FFRs do seem to represent some categorical information under perceptually demanding listening tasks involving contextual processing or noise degradation (Carter & Bidelman, 2023; Rizzi & Bidelman, 2023). It is possible the more gradient FFR representations observed here are due to the relatively low demands of our phonetic categorization task. Our use of RSA allowed us to directly compare how closely neural responses resembled both gradient and categorical information, which prior work did not address (Carter & Bidelman, 2023; Rizzi & Bidelman, 2023). That FFRs reflected gradient stimulus features does not necessarily contradict these prior findings of categorical representations in brainstem responses. Rather our data reinforce the notion that gradient information is weighted more heavily in subcortical speech coding.

While ERP-derived RDMs represented more categorical information and less gradient information than FFRs, they represented slightly more gradient than categorical information overall. This finding is consistent with prior work demonstrating that cortical responses represent both gradient and categorical stimulus information dependent on the time course of speech coding. Bidelman et al. (2013) found that cortical RDMs pattern categorically ∼200 ms after speech onset. However, gradient representations have been shown to persist beyond the obligatory waves of auditory cortical ERPs (i.e., P1-N1-P2), lasting at least 900 ms in cortical response activity (Sarrett et al., 2020; Toscano et al., 2010). Using RDMs from MEG recordings, Beach et al. (2021) found both phonemic and subphonemic information were represented cortically with subphonemic representations lasting longer (∼600 ms) during an active compared to passive CV categorization task. While we did not parse the temporal dynamics of speech processing (e.g., time-varying RDMs), we similarly find that over ∼1000 ms after speech onset, cortical activity represents both coding schemes and carries slightly more gradient than categorical information. This differential weighting likely reflects the time course of cortical speech coding as categorical representations emerge later in time (Bidelman et al., 2013). Here, the comparison of gradient vs. categorical presentation across distinct levels of the central auditory pathway is novel. In this regard, we find that both brainstem and cortical representations carry gradient speech cues, but that ERP representational geometry shows a further expansion toward categorical information that is not observed subcortically. That subphonemic (gradient) representations are maintained across this time course challenges early models of categorical perception that within-category information is discarded in favor of maintaining only between-category information (Liberman et al., 1957). We find a stronger weighting of gradient (within-category) relative to categorical (between-category) features in both subcortical and cortical responses, suggesting subphonemic detail is preserved in auditory encoding across the ascending auditory system.

### 4.5 Neural weighting of gradient vs. categorical information did not predict behavior

Our last question pertained to whether the pattern of brainstem and cortical speech representations account for gradient vs. discrete listening strategies at the perceptual level (Beach et al., 2021; Bidelman et al., 2013; Chang et al., 2010; Ou & Yu, 2022). Using similar RDM methods applied to scalp-level (sensor) FFRs/ERPs elicited by a VOT continuum, Ou and Yu (2022) found more discrete listeners had decreased similarity between subcortical and cortical representations, suggesting a larger transformation of neural representations away from stimulus acoustics to represent categorical information. However, they assessed perceptual gradience using a 2AFC task, which tempers the interpretation of their findings since gradience and consistency are confounded in a binary forced-choice paradigm (Apfelbaum et al., 2022; Kapnoula et al., 2017; McMurray, 2022). In other words, Ou and Yu (2022) may have been assessing neural correlates of perceptual consistency rather than gradience/categoricity, per se.

Using a VAS task to tease apart these perceptual factors, we did not observe a relationship between RSA measures and individual differences in categorization behavior. This suggests categorization consistency and gradience, at the perceptual level, do not depend on how closely electrophysiological responses resemble categorical and gradient representations. It is likely that individual differences in categorization decisions arise outside the bounds of the auditory brainstem-cortical pathways, e.g., from higher-level cognitive processing and frontal language regions (Bidelman et al., 2021; Binder et al., 2004; Carter et al., 2022; Fuhrmeister & Myers, 2021) or activity in supramarginal gyrus which has been linked to category decisions (Al-Fahad et al., 2020; Mahmud et al., 2021). Our primary focus in the current study was to assess how two primary stages of the central auditory system (midbrain, primary auditory cortex) map to categorization behaviors. This focus prompted us to analyze source activity rather than electrode-level activity which provides a purer measurement of the output from different neural generators. Thus, though they clearly reflect perceptually-relevant factors in speech categorization, it is possible that the auditory-centric RDMs we constructed from EEG activity in source space are too distal to perception. Collectively, our RDM findings suggest that while nascent categorical representations exist at the brainstem level, the most salient abstraction likely occurs in higher auditory-linguistic brain regions. This notion is consistent with many fMRI studies that show phonetic categories are strongly represented in frontal language regions (Blumstein et al., 2005; Lee et al., 2012; Luthra et al., 2019; Myers, 2007; Myers et al., 2009).

## 5. Conclusion

Collectively, findings from this study strengthen recent work linking individual differences in categorization to other perceptual abilities, namely SIN perception. Our findings shed new light on the neurophysiological bases of categorization consistency, suggesting consistent speech coding at either auditory brainstem or cortical levels (both not both concurrently) facilitates consistent auditory perception and phoneme labeling. Furthermore, our RSA findings provide evidence that brainstem FFRs reflect more gradient acoustic features compared to cortical ERPs, while ERPs reflect more categorical information than FFRs.

## Acknowledgements

This work was supported by the NIH/NIDCD R01DC016267 (awarded to G. M. B.) and F31DC023124 (awarded to R. R.). The authors thank Tessa Bent, Jennifer Lentz, and Samantha Gustafson for their comments on earlier versions of this manuscript and Elaina Lewis for assistance in recruitment and data collection.

## Author contributions

R. R. and G.M.B. designed the experiment, R.R., J.R.S., and Z.E. collected the data, R.R. and G.M.B. analyzed the data, and all authors wrote the paper.

## Data Availability

Data will be made available from the corresponding author upon reasonable request.

## Footnotes

1 Due to data quality issues, we excluded 3 listeners’ EEGs, resulting in 37 listeners included in the RDM analysis.

